# An automated homecage system for multiwhisker detection and discrimination learning in mice

**DOI:** 10.1101/2020.04.27.063750

**Authors:** Sarah M. Bernhard, Jiseok Lee, Mo Zhu, Alex Hsu, Andrew Erskine, Samuel A. Hires, Alison L. Barth

## Abstract

Automated, homecage behavioral training for rodents has many advantages: it is low stress, requires little interaction with the experimenter, and can be easily manipulated to adapt to different experimental condition. We have developed an inexpensive, Arduino-based, homecage training apparatus for sensory association training in freely-moving mice using multiwhisker air current stimulation coupled to a water reward. Animals learn this task readily, within 1-2 days of training, and performance progressively improves with training. We examined the parameters that regulate task acquisition using different stimulus intensities, directions, and reward valence. Learning was assessed by comparing anticipatory licking for the stimulus compared to the no-stimulus (blank) trials. At high stimulus intensities (>9 psi), animals showed markedly less participation in the task. Conversely, very weak air current intensities (1-2 psi) were not sufficient to generate rapid learning behavior. At intermediate stimulus intensities (5-6 psi), a majority of mice learned that the multiwhisker stimulus predicted the water reward after 24-48 hrs of training. Both exposure to isoflurane and lack of whiskers decreased animals’ ability to learn the task. Perceptual learning was assessed and following training at an intermediate stimulus intensity, perception was likely heightened as mice were able to transfer learning behavior when exposed to the lower stimulus intensity. Mice learned to discriminate between two directions of stimulation rapidly and accurately, even when the angular distance between the stimuli was <15 degrees. Switching the reward to a more desirable reward, aspartame, had little effect on learning trajectory. Our results show that a tactile association task in an automated homecage environment can be monitored by anticipatory licking to reveal rapid and progressive behavioral change. These Arduino-based, automated mouse cages enable high-throughput training that facilitate analysis of large numbers of genetically modified mice with targeted manipulations of neural activity.

## Introduction

The whisker system has been extensively used in mice and rats to study the organization and response properties of neurons in the somatosensory system. The barrel cortex, a precise somatotopic map of identified facial vibrissae in the neocortex, facilitates the targeted analysis of whisker-dependent stimulus response properties and experience-dependent plasticity. Stimulation of a single whisker has been used to map receptive field properties of cortical neurons (1,2), as well as drive experience-dependent plasticity (3–6). Indeed, with intensive training, mice and rats can use a single whisker to detect object location (1,7,8), indicating that individual whisker activation can be behaviorally meaningful.

Because the whiskers are typically used together during normal sensory activation, multiwhisker stimulation has increasingly been used to study the response transformations and plasticity of cortical neurons (9–11). New studies show that multiwhisker stimuli can potently activate cortical neurons in ways that were not predicted by single-whisker stimuli (10,12). In addition, vibrissae can be used not only for active sensation, reflected in whisking behavior that often accompanies exploration of novel objects, but also for detection of low-frequency input from the environment. For example, harbor seals can track a decoy through water by tracking alterations in local currents, a task that is whisker-dependent (13).

Because multiwhisker stimuli are an ethologically appropriate way to activate the facial vibrissae, we reasoned that these stimuli might be an excellent probe to investigate learning and plasticity in mice. Indeed, we have recently shown that multiwhisker stimuli are readily detected by mice and can be used in a sensory learning task that drives plasticity in cortical circuits (11). Here we sought to determine how multiwhisker stimuli, delivered through a gentle air current directed at the large facial vibrissae of mice, could be used to drive learning behavior in an automated sensory association task. These stimuli are quantitatively different from those used as punishment in other investigations which use airpuff intensities that are 5-100x greater than those deployed in our studies and are often directed toward the animal’s face or eye. In contrast, the stimuli used here were of low intensity and specifically targeted at the distal ends of the large facial vibrissae.

We examined the parameters required for mice to learn how to detect and discriminate multiwhisker deflections caused by an air current directed to the large vibrissae. Establishing this stimulus training paradigm in rodents would be useful for neurobiological studies, as it can be adapted to homecage training in freely-moving animals and is well-suited for cellular analysis of cortical circuits, since the anatomical region corresponding to the stimulated whiskers is broad and experimental analysis does not need to be targeted to a single barrel column. Furthermore, automation of the behavioral set up allows for an increase in throughput, with minimal interaction with the experimenter and less variability in training conditions. Our results show that mice rapidly learn to associate a multiwhisker stimulus with a reward, that they show an exquisite sensitivity to discriminate different directions of stimulation, and that sensory association training (SAT) reduces perceptual thresholds for stimulus detection.

## Materials and methods

### Animals

Behavioral data was collected from 131 C57/BL6 mice (Harlan Laboratories); ages ranged from postnatal day 22 (P22) – P28. Mice were housed individually during training. Animals were exposed to a 12-hour light-dark cycle schedule with lights on at 7am and had free access to food and water, the only source of which was dispensed from a recessed lickport in the custom-built chamber. Animals were given at least 24 hours to acclimate to the cage before SAT, during which there was no sensory stimulus coupled to water delivery. Approximately 1-3 ml of water was dispensed each day. All experiments conducted were approved by Carnegie Mellon University Animal Care and Use Committee.

### Stimulus calibration

Throughout SAT, stimulus intensity was set to a constant level using a gas regulator (Fisherbrand). To ensure accurate calibration of stimulus intensity, a pressure transducer (NXP USA Inc.) was used to provide an exact measurement of pressure at the opening of the air tube. Actual air pressure at the whiskers was lower than at the air tube opening, located ~4 cm above the whiskers (Fig 1), and because animals self-positioned at the nosepoke, it was not possible to determine small variations in the specific whiskers activated during training. Three different stimulus intensities were used in these studies: 1-2 psi (abbreviated as 2 psi), 5-6 psi (abbreviated as 6 psi), or 9 psi air puffs. We calculated that a 6 psi stimulus was equivalent to 0.4 bar. Because 1 psi pressure intensity was difficult to control with a conventional gas regulator, a second miniature gas regulator (PneumaticPlus) was used in series. The same training paradigm for acclimation (24 hrs) and training days (48-72 hrs) was used for all stimulus intensities.

**Figure 1.**
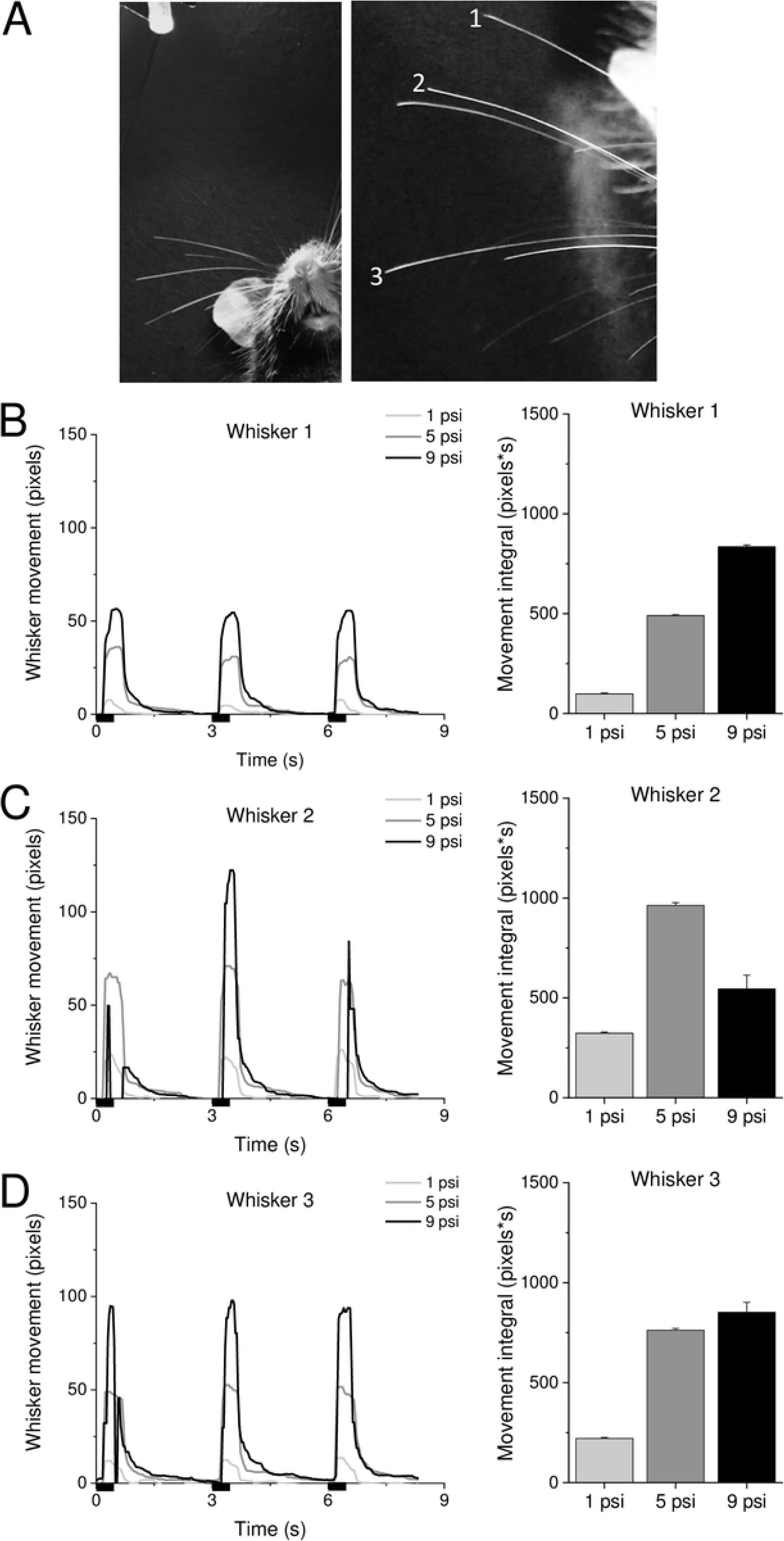
Airpuff intensity and whisker movement. A) Left, air nozzle tip is located approximately 4 cm above whiskers. Right, three whiskers analyzed in B-D. B) Left, example movement traces of the whisker indicated in (A) at 3 different air puff strengths; 1, 5, and 9 psi. Black bars indicate air puff duration (500 ms). Right, average movement (area under the curve in the left graph) of the whisker indicated in (A) at 3 different air puff strengths. Average of 50 air puffs. C-D) Same as in B, for whiskers 2 and 3, respectively.

### Whisker movement analysis

To calibrate evoked whisker movements, air current stimulation was delivered to an anaesthetized mouse mounted with an air nozzle ~4 cm above and to the right of the animal’s whiskers. Whisker movement was video recorded while receiving air current stimuli at 1, 5, or 9 psi. Stimuli were delivered 50 times for each intensity, every 3 seconds for 500 ms (Fig 1). Movement of the A3 whisker was tracked using a variant of DeepLabCut ((14), https://github.com/RoboDoig/dlc-cloudml) trained to identify the whisker tip position. Displacement of the whisker was analyzed from the DeepLabCut model using a custom MATLAB script.

### Sensory Training Paradigm

Automated homecage training chambers for singly-housed animals were custom-made at Carnegie Mellon University. They consisted of a standard 7×12” mouse cage with a custom-built 3×5” stimulus chamber attached that contained a recessed lick port, 1/16” in diameter, which was fixed 2 cm above the base of the chamber ((11); Fig 2). Air currents were delivered ~4 cm above and 2.5 cm to the right of the recessed lick port, to ensure that they were directed at the distal tips of the whiskers. The infrared (IR) beam (Adafruit; Table 1) was also recessed and located approximately 1 cm in front of the lick port to signify whether a nose poke had occurred. To record licking behavior, a capacitive touch sensor (Adafruit; Table 1) was attached to the metallic lick port and a lick was recorded when the capacitance reached threshold. Data output from the lick sensor and IR beam was updated every 100 ms. Importantly, this design does not detect individual licks, which might occur ≥10 Hz. Furthermore, any licks that occurred at any point within the 100ms period were counted as one lick.

**Figure 2.**
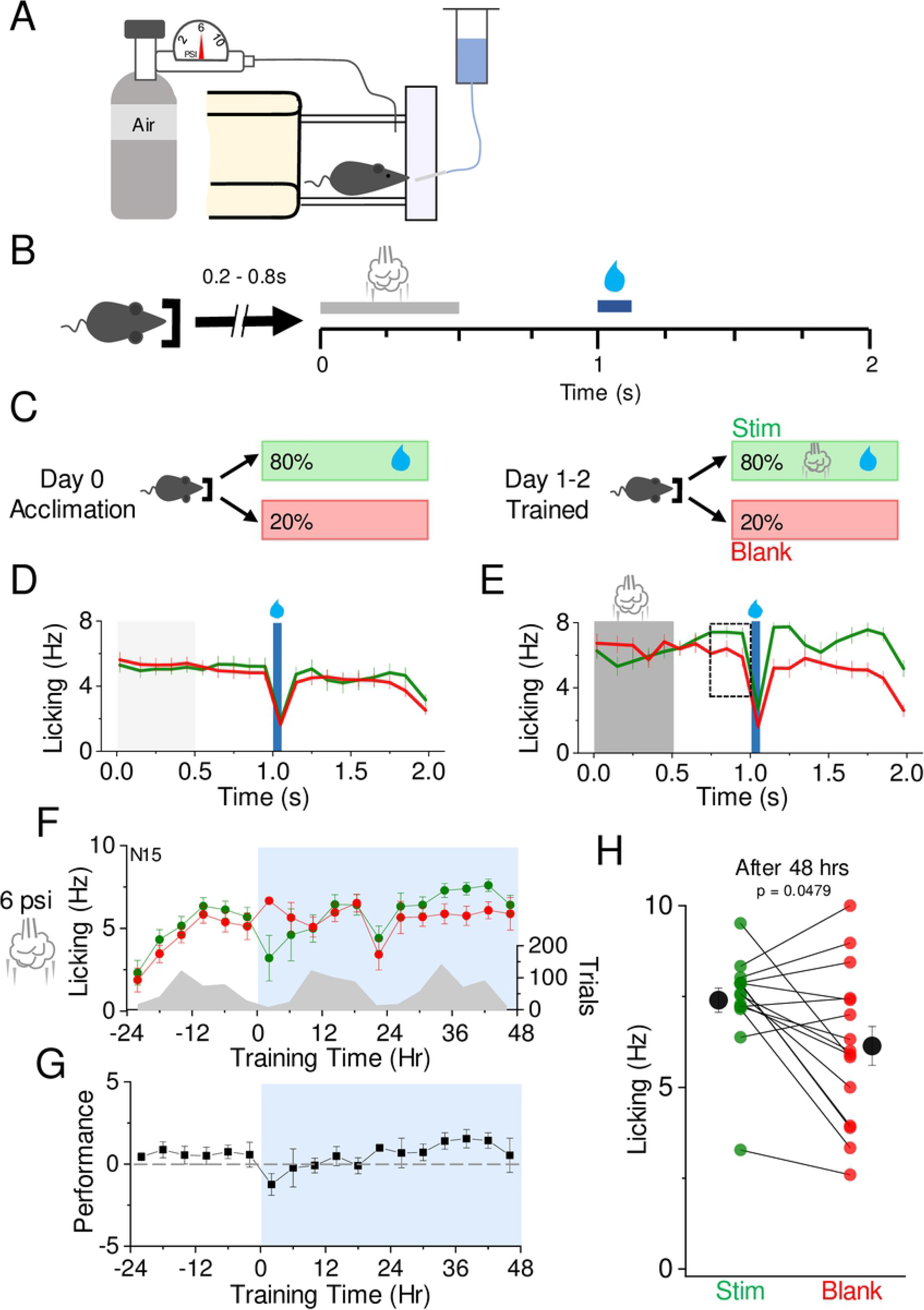
SAT drives changes in anticipatory licking. A) Left, schematic of the homecage training apparatus. Right, freely-moving mouse positioned at lick port. B) Behavioral paradigm for association of air puff and water delivery. Animals initiate trials by breaking an infrared beam at the nosepoke, resulting in a random delay ranging from 0.2-0.8 s followed by a 500 ms air puff (grey bar). Water delivery occurs 500 ms after the end of the air puff, lasting 75 ms. Trials cannot be reinitiated for 2 s following air puff onset. C) Reward contingencies during training. Left, during the initial 24 hour acclimation period, animals receive water on 80% of initiated trials with no air puff. Right, during the training period, animals receive air puff and water on 80% of initiated trials. D) Mean lick frequency for water delivery (green) or blank (red) trials for the acclimation period, binned at 10Hz. Water delivery time indicated by a blue bar. E) As in (D) but with air puff-water coupling. Air puff timing indicated by grey shading. F) Mean lick frequency for water and blank trials, binned at 4 hr intervals. Air puff set at 6 psi and association training is indicated at t=0 (12 noon/daylight period). Mean lick frequency for water delivery (green) or blank (red) trials is overlaid upon mean number of initiated trials (grey) across training days. G) Mean performance (lick frequency for water trials – lick frequency for blank trials) for each 4 hour bin during the acclimation period (−24 to 0 hr) and training phases (0 to 48 hrs). H) Mean lick frequency for the last 20% of total trials for each animal exposed to 6 psi intensity air puff. N=15 animals.

**Table 1.** Key resources for behavioral chambers.

| Product name | Company | Product ID |
| --- | --- | --- |
| Leonardo | Arduino | A000057 |
| Yún Shield v2.4 | Dragino | N/A |
| Relay Shield for Arduino v2.1 | DFRobot | DFR0144 |
| Standalone Momentary Capacitive Touch Sensor | Adafruit | 1374 |
| IR Break Beam Sensor | Adafruit | 2167 |
| Solenoid Valve | The Lee Company | LHDA1233115H |
| Gas regulator | Fisherbrand | 10-575-105 |
| Miniature Air Pressure<br>Regulator | PneumaticPlus | PPR2-N02BG-2 |
| Pressure Transducer | NXP USA Inc. | MPX5100GP |

Trials were self-initiated by an IR beam-break at the nosepoke entry port in the stimulus chamber (Fig 2). Once a trial started, there was a random delay ranging from 200-800 ms before stimulus delivery, to ensure that the sensory association would be made to the stimulus and not to the operant cue from the nosepoke. This random delay was followed by 500ms of the air puff stimulus. If the mouse was in the acclimation period, then no air puff would be delivered during this time. After a 500 ms break, water was delivered for 75 ms, equating to approximately 15 μL. There was a 925 ms break following water delivery, during which the next trial could not be initiated. A relay shield (DFRobot; Table 1) was used to activate solenoids (The Lee Company; Table 1) at precise times during the trial for both stimulus and water delivery. To disguise possible auditory cues promoted by the relay shield, non-stimulus trials activated separate relays that did not gate any air current.

An Arduino Leonardo was used to run and maintain the paradigm. The Yún Shield (Dragino; Table 1) connected the set up to the local Wi-Fi router and stored data collected from experiments. This device uploaded real time data to the internet for remote access.

### Isoflurane exposure

For experiments to test the effects of inhalation anesthetics on sensory association learning, mice were exposed to isoflurane anesthesia in an enclosed glass jar for approximately 30-60 s until the hindlimb withdrawal reflex was absent. Anesthetic exposure occurred only once at midday prior to the first day of acclimation. Isoflurane-exposed animals were then housed in the training chamber for a 24 hr acclimation period followed by 48 hrs of sensory association training using a 6 psi airpuff stimulus.

### Whisker removal

At some low incidence in standard animal housing, mice will spontaneously barber the whiskers of cagemates so that no large facial vibrissae remain; indeed, barbered animals typically have both fur and vibrissae removed. We took advantage of this natural behavior and used barbered mice to investigate whether whiskers were required for association learning in this automated set-up, without anesthetic confounds. Barbered mice were exposed to the sensory association task using a 6 psi airpuff stimulus as described above.

### Direction discrimination

Discrimination learning was tested by using two different oriented air puffs with the same paradigm as described for sensory association training above, where 80% of trials used one direction and were coupled to the water reward and 20% of trials were at a different direction and were unrewarded. The two air puffs were delivered in a cylindrical association chamber with a central platform that the animal used to approach the nosepoke for trial initiation. Air tubes were oriented around this cylinder so that the delivery angle could be precisely controlled. The platform contained a cut out on the right side, in the location air puffs were delivered, to ensure that air puffs could be administered below the animal for upward deflection of the whiskers.

A second solenoid was used to deliver the unrewarded air puff. Unrewarded directional airpuffs (500 ms) were also delivered according to the same trials parameters as described above: initiated by a nosepoke, with a random delay before presentation and no coupled water reward.

### Altered reward contingency

Reward contingency could be digitally adjusted using the source code to alter stimulus-reward frequency. To determine whether reducing the frequency of reward trials would influence learning trajectories, the percentage of trials coupled to water was adjusted from 80% to 50% of initiated trials, and the remaining 50% of trials were blank trials. Airpuff intensities were set at 6 psi and mice received the standard 24 hr acclimation and 48-72 hr training period.

### Perceptual learning air puff intensity during training

Mice were exposed to the standard acclimation day and two days of 6 psi training. Following these two days of training at 6 psi, mice received another day of training with the air puff intensity decreased to 1 psi. Because the larger gas regulator is not as precise at producing air puffs with 1 psi intensity, the smaller gas regulator was used. A pressure transducer confirmed that the pressure was exact.

### Aspartame training

To examine the effect of enhanced reward in SAT, we calibrated drinking preference to aspartame, sucralose, saccharine, and sucrose. Animals showed a modest preference for aspartame compared to other sweeteners and so this was used for subsequent experiments. When provided with either 10% aspartame or water as their sole source of hydration, mice showed a marked preference for aspartame; thus, we used aspartame in place of water to enhance reward valence. Animals were acclimated to the training cage with water provided through the lickport. After acclimation, water was replaced with 10% aspartame and the same 80%-stimulus/reward, 20% blank trial schedule was introduced. Anticipatory licking was calculated as described above. For aspartame-trained animals, aspartame solution consumption was modestly higher than in water-reward trials (~4.5 mls aspartame solution versus ~3 mls for water).

### Behavioral analysis

Behavioral data obtained from experiments was analyzed using custom scripts in MATLAB (https://github.com/barthlab/Sensory-association-training-behavior). All licks times were adjusted to the beginning of the trial at air puff onset, following random delay. This readjustment was necessary to be able to compare lick times across trials with different random delay times. Licks were counted if they had taken place in the 700 – 1000 ms time window after the random delay, which was 300 ms directly before water delivery. Only these licks were analyzed to discriminate between anticipatory and consummatory licks. Anticipatory licks were separated based on stimulus and blank trials and binned into 4 hour intervals. The values were then converted into Hz. Performance was calculated by subtracting the lick rate of blank trials (Lick blank; L_b_) from water-rewarded trials (Lick water; L_w_) for each 4 hour time bin (performance=L_w_−L_b_). The last 20% of trials were analyzed and the lick rate for water trials was compared to blank trials. Behavior analysis was conducted for each animal and then averaged with other animals in the same experiment.

### Statistical analysis

A Wilcoxin rank sum test was carried out to evaluate absolute differences in licking in stimulus (L_w_) versus blank (L_b_) for the last 20% of trials after 48 hrs of SAT for animals within an experimental group, to determine whether specific training conditions were sufficient to alter behavior. The whisker-dependent sensory association behavioral paradigm developed here was easily adapted to a variety of different stimulus and reward conditions. Although in theory statistical comparisons could be made across experimental groups to identify optimal parameters for training, in practice wide variation in animal behavior during early learning - even within experimental groups- made it difficult to identify statistically significant differences. Despite the large number of animals in different test groups, experiments were generally underpowered to detect small differences in performance across conditions after 48 hrs of training. Thus, we did not directly compare behavioral changes across training conditions.

## Results

### Stimulus-evoked whisker movement

We first calibrated the degree of whisker deflection introduced by the gated air current, using video analysis in a head-fixed, anaesthetized mouse (Fig 1; S1-3 Videos). The air current was gated by a solenoid valve, and the position of the tube relative to the vibrissae was similar to that in the homecage training apparatus, about 4 cm. Although this does not necessarily recapitulate the stimulus in a freely-moving animal with variable positioning across trials, it enabled us to determine the maximal effects of different stimulus levels across multiple whiskers, i.e. those closest and further from the stimulus source. Individual whiskers were identified and movement tracked using custom software ((14), https://github.com/RoboDoig/dlc-cloudml). For whiskers closer to the stimulus, whisker movement was continuous when the solenoid valve was open (500 ms) and scaled with stimulus intensity (Fig 1B). More distant whiskers showed lower deflections that did not necessarily scale, likely because of non-linearities in how air currents disperse in a complex environment, as well as variations in the length of individual whiskers (Fig 1C,D). Overall, we found that the positioning of the air current above and to the right of the animal’s head lead to a broad and prolonged movement of ~1-5 mm for all the whiskers depending on location. Because animals are freely-moving and because positioning will differ across individual trials, it is likely that this controlled environment does not precisely reflect stimulus properties experienced by animals during sensory association training. However, these measurements provide a reference point for other studies that might employ an air current stimulus.

### Automated training for sensory association learning

Our prior studies have used a 6 psi multiwhisker stimulus coupled to a water reward to drive association learning (11). Because animals were not water deprived and could freely initiate trials, this training environment has the advantage of being both low-stress for the animal and scalable so that multiple animals can be trained in parallel with little intervention from the investigator.

Individually-housed animals were acclimated to the training cage for 24 hrs prior to sensory association training (SAT; Fig 2A-C). After 24 hrs, we introduced the airpuff stimulus, so that animals only received water when it was coupled to a prior airpuff, for 80% of initiated trials. Blank trials occurred at the remaining 20% of trials, when the animal initiated a nosepoke but neither stimulus nor water were delivered. This trial structure allowed us to compare licking for stimulus versus blank trials for each animal to obtain an individual metric that reflects learning for each animal. SAT-dependent changes in anticipatory licking 300 ms prior to water delivery was used as an indicator of sensory association learning (Fig 2D,E), which progressively increased in stimulus-reward trials over the training period. At the onset of training, animals displayed a suppression of licking behavior during stimulus trials reflected in greater licking on “blank” trials compared to stimulus trials, likely due to the novelty of the stimulus (* in Fig 2G). Animals rapidly habituated to the stimulus, and analysis of performance (L_w_-L_b_) shows a steady increase in anticipatory licking over the 48 hr period of SAT (Fig 2F,G). Increases in performance were driven primarily by increased licking in stimulus trials, not suppressed licking in “blank” trials.

Although licking behavior was variable across animals, the majority (11/15) of animals showed an increase in anticipatory licking by the end of 48 hrs of SAT and this change in behavior was statistically significant (Fig 2F-H). These results show that SAT rapidly drives changes in behavior, measured both by habituation to the stimulus in the first 24 hrs of training and by significant increases in anticipatory licking in the majority of animals after 48 hrs of training.

### SAT is whisker-dependent

To determine whether animals were using the facial vibrissae for sensory association learning, we tested whether they could perform the task in the absence of the large facial vibrissae. Initially, we carried out control experiments on mice exposed to isoflurane anesthesia where whiskers were not removed, since this is typically used to immobilize animals for whisker plucking and would be required for comparison. Inhalation isoflurane exposure was brief (~1 minute) and was carried out prior to the first acclimation day in the training cage (Fig 3A). Surprisingly, isoflurane exposure alone, where all whiskers were intact, was sufficient to suppress SAT-associated changes in behavior after 48 hrs of SAT with the 6 psi stimulus (Fig 3A-C). An increase in anticipatory licking was almost never observed in the isoflurane-exposed training cohort (only 1/6 showed greater L_w_-L_b_ after 48 hrs of SAT). Because animals showed a transient decline in licking to stimulus trials in the first few hours of training, it appears that they may be able to initially detect the stimulus. Thus, we conclude that isoflurane exposure may suppress rapid learning in SAT.

**Figure 3.**
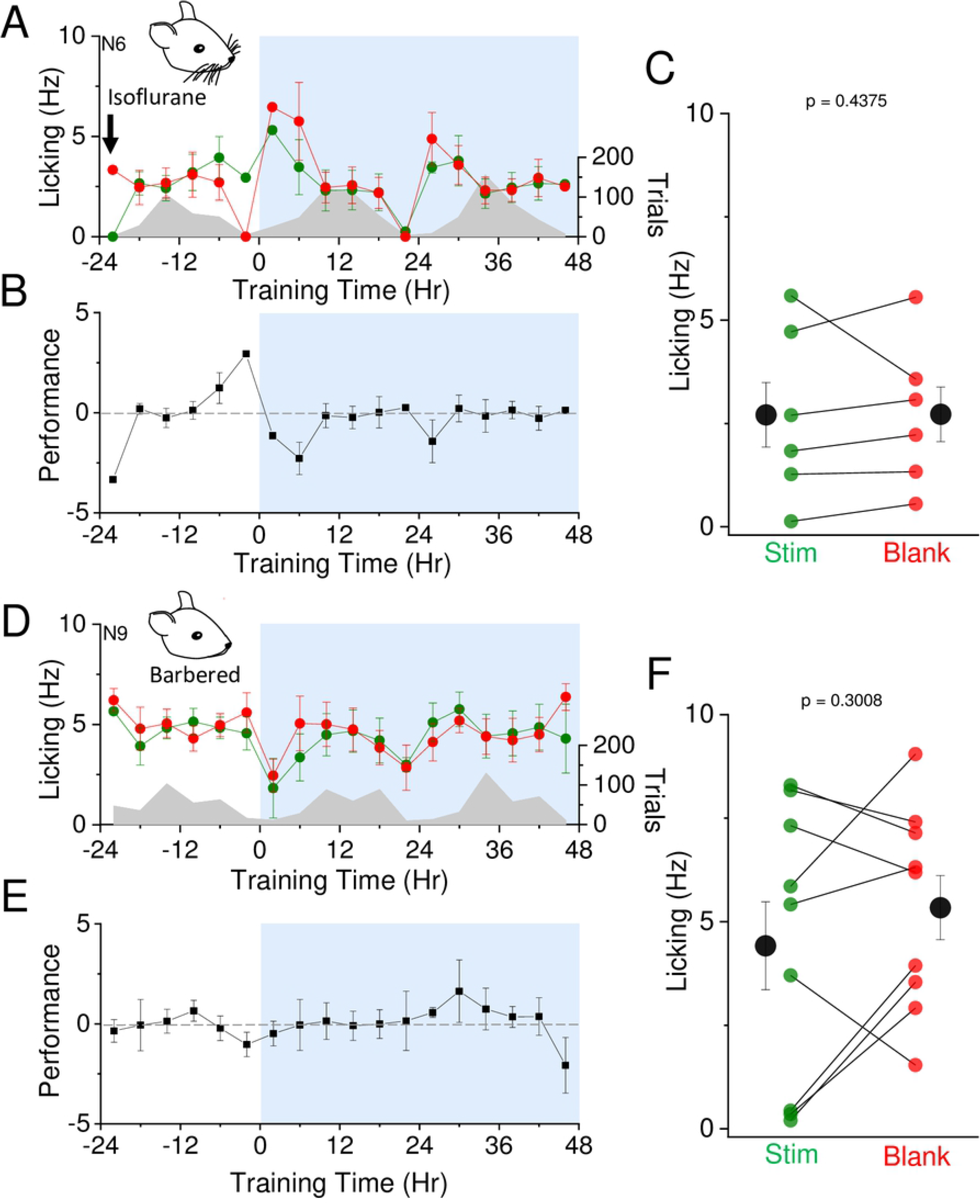
Isoflurane and absence of whiskers suppresses learning. A) Mice were exposed to isoflurane until breathing slowed and then placed in training chamber for acclimation period followed by training. Air puff set at 6 psi and association training is indicated at t=0 (12 noon/daylight period). Mean lick frequency for water delivery (green) or blank (red) trials, binned at 4 hr intervals, is overlaid upon mean number of initiated trials (grey) across training days. B) Mean performance (lick frequency for water trials – lick frequency for blank trials) for each 4 hour bin during the acclimation period (−24 to 0 hr) and training phases (0 to 48 hrs). C) Mean lick frequency for the last 20% of total trials for each isoflurane induced animal exposed to 6 psi intensity air puff. N=6 animals. D) Whiskers were barbered by cage mates prior to training. No isoflurane was used. Same as in (A) but with barbered animals. E) Same as in (B) but with barbered animals. F) Same as in (C) but with barbered animals. N=9 animals.

To determine whether facial whiskers were required for sensory learning, we used an alternate approach, taking advantage of a natural behavior in laboratory mice, where the large facial vibrissae are sometimes removed by cagemates. We opportunistically identified animals aged P22-28 from our C57/BL6 colony that lacked whiskers and tested them with the SAT task. On average, barbered animals failed to increase anticipatory licking after 48 hrs of SAT. It is possible that barbered animals retained some fine vibrissae at the mystacial whisker pad, or they could detect the air current using other whiskers that remained (for example, around the eyes, at the ears, or around the mouth). Lack of the transient decline in licking behavior during stimulus trials suggests that recognition of the air puff is hindered without these large facial vibrissae. These data suggest that the large facial vibrissae are required for learning in this sensory association task.

### Stimulus intensity influences learning

Our prior studies used a moderate stimulus intensity that balanced animal participation and learning speed. To systematically determine how stimulus intensity would influence the trajectory of behavioral change, we compared the effects of training using a lower and a higher-intensity airpuff stimulus. When the airpuff stimulus was low (2 psi), 48 hrs of SAT was not sufficient to drive a significant change in anticipatory licking on average across the test group. The lack of significance was driven primarily by heterogeneity in comparative lick frequency, since some animals showed a large increase in anticipatory licking and others showed no difference or greater licking on “blank” trials (12/21 animals showed L_w_>L_b_; Fig 4C). In contrast, SAT with higher-intensity (9 psi) airpuff stimuli did drive significant change after 48 hrs SAT on average (6/7 animals showed L_w_>L_b_; Fig 4D-F).

**Figure 4.**
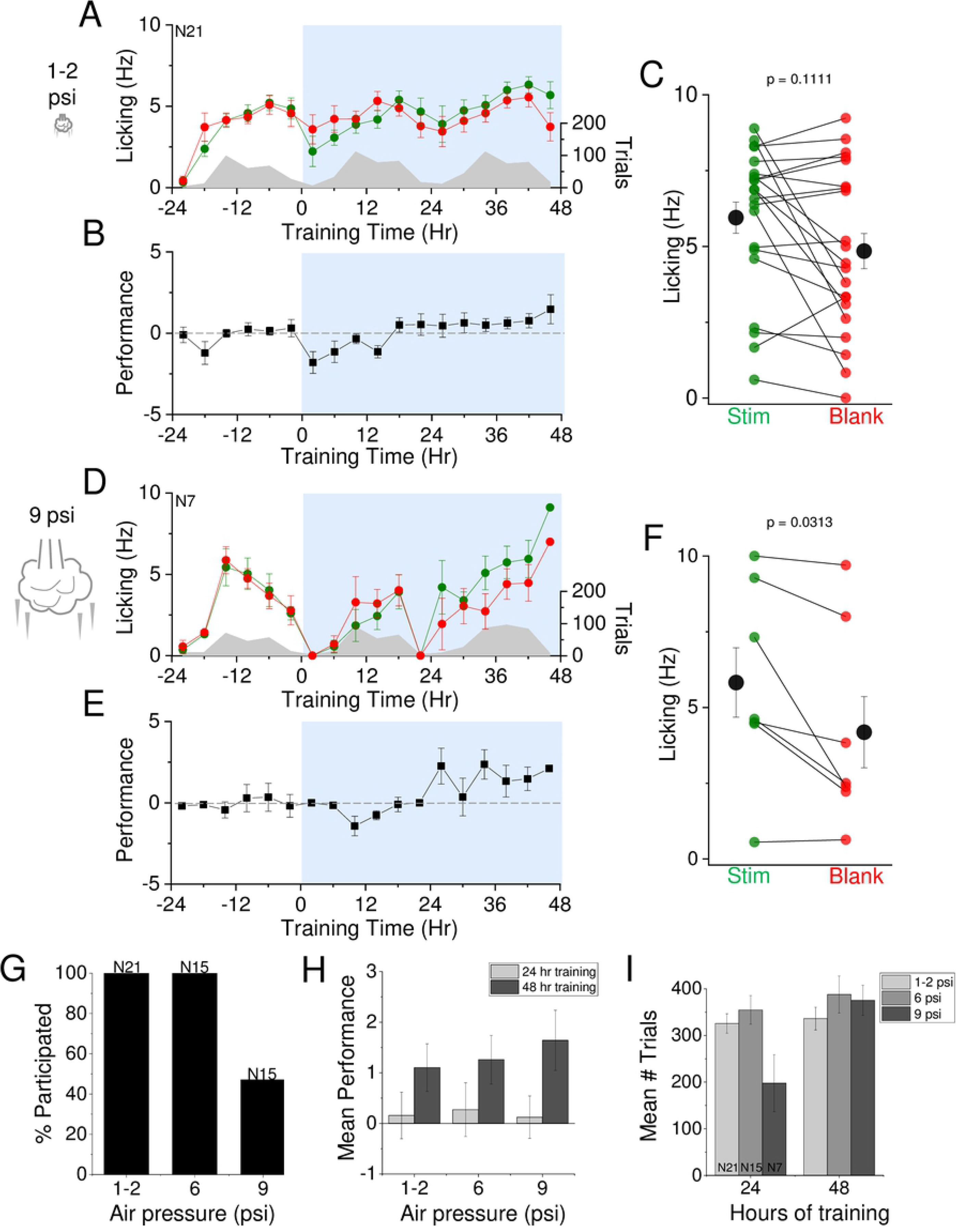
Stimulus intensity alters learning trajectory. A) Air puff was set at 1-2 psi and association training is indicated at t=0 (12 noon/daylight period). Mean lick frequency for water delivery (green) or blank (red) trials, binned at 4 hr intervals, is overlaid upon mean number of initiated trials (grey) across training days. B) Mean performance (lick frequency for water trials – lick frequency for blank trials) at 1-2 psi air puff intensity for each 4 hour bin during the acclimation period (−24 to 0 hr) and training phases (0 to 48 hrs). C) Mean lick frequency for the last 20% of total trials for each animal exposed to 1-2 psi air puff intensity. N=21 animals. D) Air puff set at 9 psi. Same as in (A) but with air puff intensity set at 9 psi. E) Same as in (B) but with air puff intensity set at 9 psi. F) Same as in (C) but with air puff intensity set at 9 psi. G) Percent participation of animals in behavioral task at different air puff intensities. H) Mean performance during the last 20% of total trials at different air puff intensities after 24 hours (light grey) and 48 hours (dark grey) of training. I) Mean number of trials for each day during the first and second days of training for different air puff intensities.

The efficacy of training with a higher-intensity stimulus were mitigated by the large number of animals that chose not to participate in the training paradigm, i.e. stopped initiating trials in the first few hours of SAT. Animal drop-out was never observed with low or medium intensity stimuli but frequently with high intensity stimuli (8/15 animals did not participate in training; Fig 4G). By 48 hrs of SAT, average performance for medium and high stimulus intensities were similar and low intensity stimuli was modestly lower. Although high stimulus intensity was correlated with a smaller number of initiated trials after 24 hrs of SAT, the mean number of initiated trials was similar across conditions by the second training day (Fig 4I). Thus, SAT with medium intensity stimuli provides a good balance between ensuring that the majority of animals participate in the training paradigm and driving rapid and significant behavioral change across the majority of participants.

### Reducing reward probability does not suppress learning

Reward probability will influence learning trajectories, since infrequent pairing of stimuli with reward can make it more difficult to build an association. We compared the trajectory of learning using a medium intensity stimulus on a modified reward schedule, where stimulus-water coupling occurred on 50% of trials, versus 80% in our initial studies (Fig 5A).

**Figure 5.**
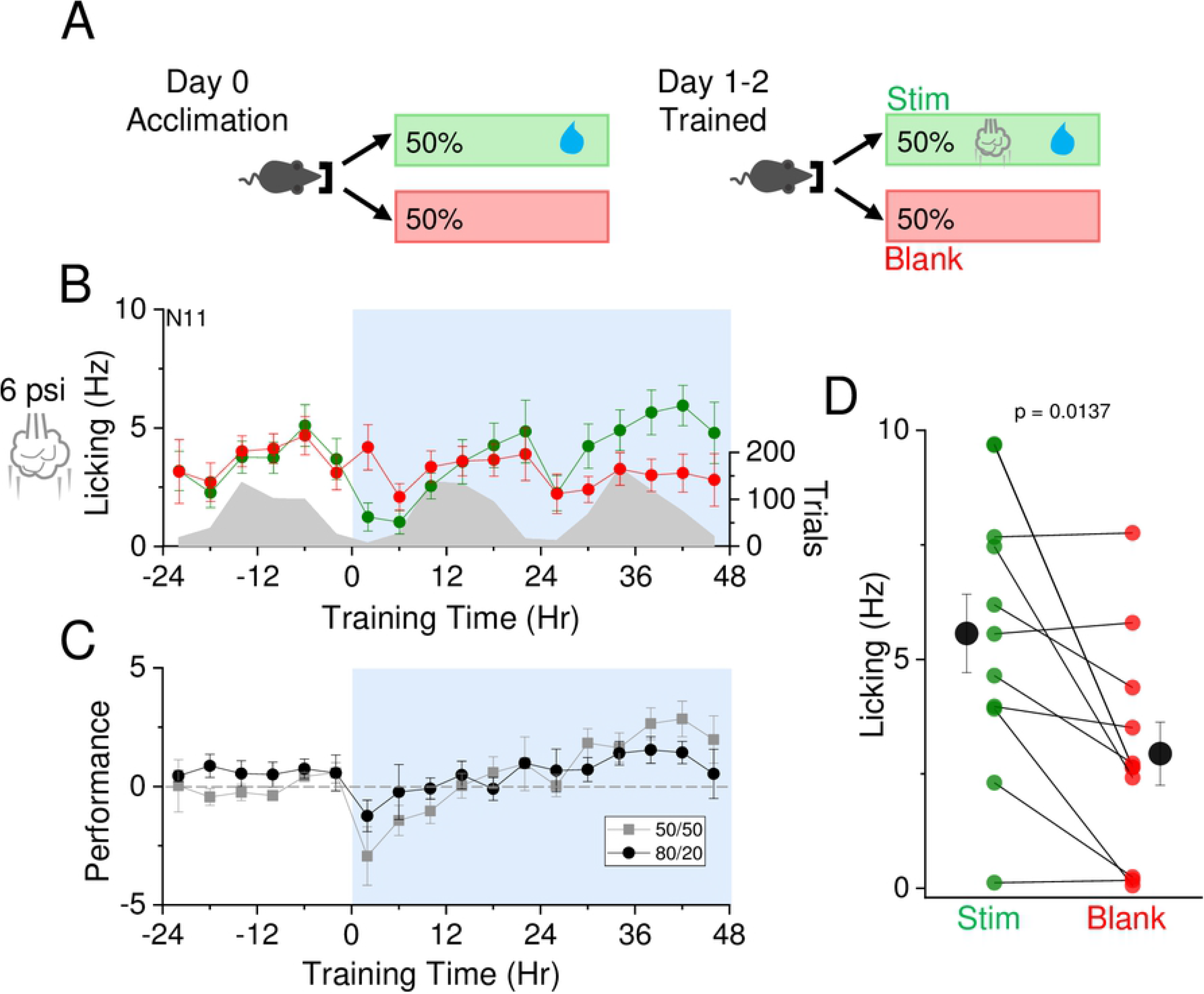
Learning is maintained with reduced reward frequency. A) Reward contingencies during training. Left, during the initial 24 hour acclimation period, animals receive water on 50% of initiated trials with no air puff. Right, during the training period, animals receive air puff and water on 50% of initiated trials, and no air puff nor water on the remaining 50%. B) Air puff set at 6 psi and association training is indicated at t=0 (12 noon/daylight period). Mean lick frequency for water delivery (green) or blank (red) trials, binned at 4 hr intervals, is overlaid upon mean number of initiated trials (grey) across training days. C) Mean performance (lick frequency for water trials – lick frequency for blank trials) of mice experiencing 50/50 paradigm (grey) or 80/20 paradigm (black) for each 4 hour bin during acclimation period (−24 to 0 hr) and training phases (0 to 48 hrs). D) Mean lick frequency for the last 20% of total trials for each animal exposed to a 50/50 contingency. N=11 animals.

Reducing the fraction of stimulus and reward trials did not slow learning trajectories over the 48 hr training period; indeed, performance was moderately enhanced using the 50% reward frequency compared to the 80% used in Figs 2 and 3. The overall number of trials conducted during 50% reward frequency was higher than with 80% reward frequency; however, the same amount of water was elicited per day. The fraction of animals that showed greater L_w_>L_b_ was similar between the two conditions (80% reward: 11/15 versus, 50% reward: 8/11; or ~72% for both). These data indicate that the automated sensory association paradigm can be modified to adjust reward contingencies at different stages of training to probe the effects on behavior.

### SAT drives perceptual learning

Prior studies have suggested that sensory stimulation can alter cortical response properties in the absence of learned associations, increasing the number of neurons that spike in response to a weak stimulus after some period of sensory exposure (15). Such a finding suggests that perceptual thresholds might be lowered in this sensory training paradigm. Perceptual learning is typically defined as long-lasting changes in perception due to practice or experience. To determine whether SAT might be associated with an increase in perceptual acuity, we trained animals using a medium-intensity stimulus (6 psi) for 48 hours, and then tested them with a low-intensity stimulus (2 psi) that by itself did not drive significant changes in behavior (Fig 6A). Because animals do not reliably show a change in licking behavior when trained for two days with a low-intensity stimulus (Fig 4A-C), increased licking responses to the low-intensity stimulus after training with medium-intensity airpuff would provide evidence for perceptual learning.

**Figure 6.**
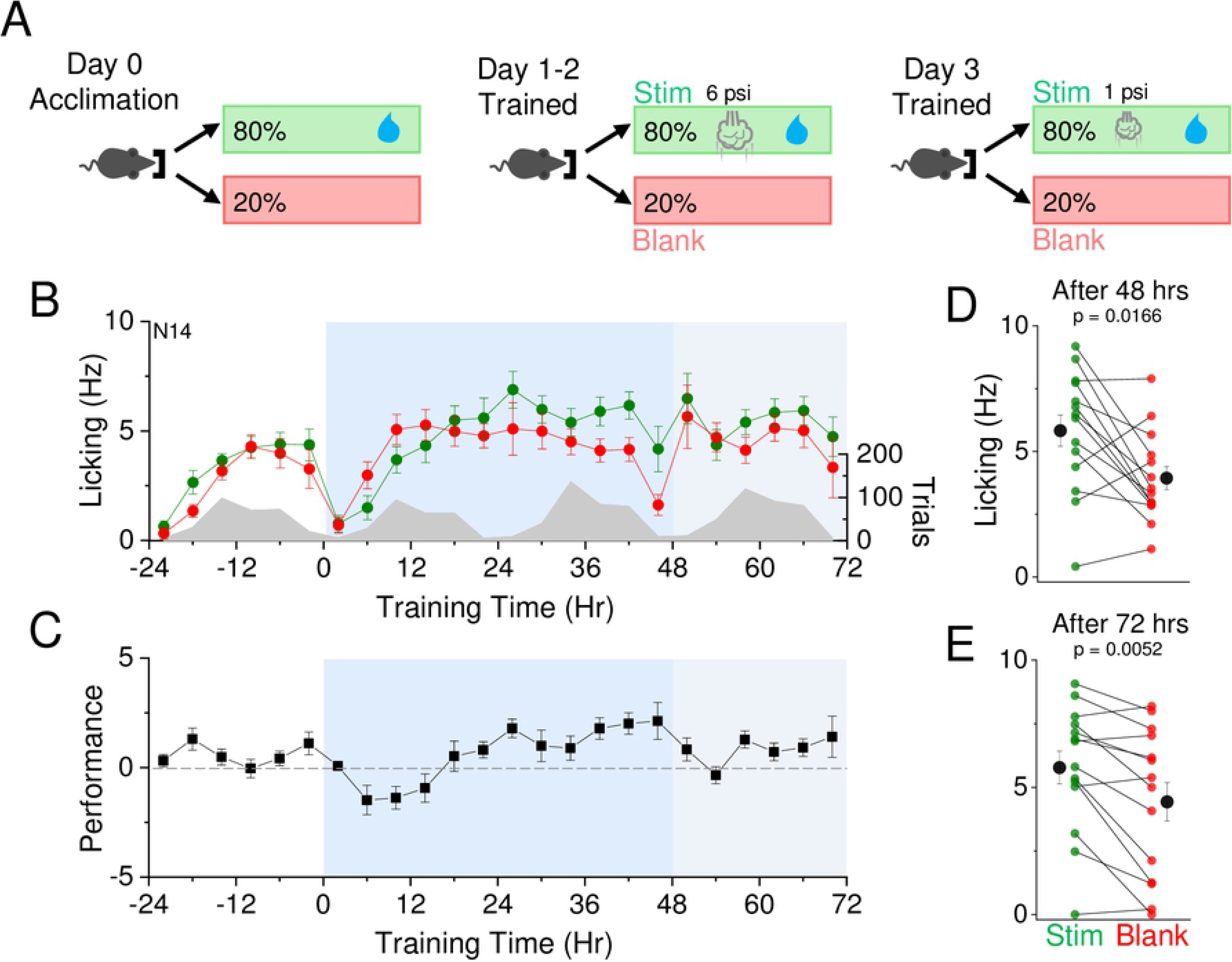
Decreased detection threshold after training suggests perceptual learning. A) Reward contingencies during training. Left, during the initial 24 hour acclimation period, animals receive water on 80% of initiated trials with no air puff. Middle, during the first training period, animals receive water and an air puff at 6 psi intensity on 80% of initiated trials for 2 days. Right, during the second training period, animals receive water and an air puff at 1 psi intensity on 80% of initiated trials for 1 day. B) Air puff association training at 6 psi is indicated at t=0 (12 noon/daylight period). Air puff association training at 1 psi is indicated at t=48 (12 noon/daylight period). Mean lick frequency for water delivery (green) or blank (red) trials, binned at 4 hr intervals, is overlaid upon mean number of initiated trials (grey) across training days. C) Mean performance (lick frequency for water trials – lick frequency for blank trials) for each 4 hour bin during the acclimation period (−24 to 0 hr) and training phases (0 to 72 hrs). D) Mean lick frequency for the last 20% of total trials for each animal after 2 days of training with 6 psi air puff intensity. E) Mean lick frequency for the last 20% of total trials for each animal after 2 days of training with 6 psi air puff intensity and 1 day of training with 1 psi air puff intensity. N=14 animals.

As expected, animals showed a significant increase in anticipatory licking after 48 hrs of training with the medium-intensity stimulus (10/14 animals showed a significant difference in licking frequency; Fig 6B-D). When the stimulus was reduced to the low-intensity airpuff at the beginning of the third training day, averaged anticipatory licking on stimulus trials was initially reduced but rapidly increased relative to “blank” trials after 24 hrs of training, a difference that was highly significant (10/14 animals showed a significant difference in licking frequency; Fig 6B,C,E). Of note, the 4 animals that did not show greater stimulus-evoked licking during low-intensity stimulus training also did not exhibit altered licking responses after 48 hrs of training at the medium-intensity stimulus. These data suggest that animals can effectively transfer the association of the medium-intensity stimulus with the water reward to a lower intensity stimulus. Thus, this training assay may be an effective and high-throughput platform to study perceptual learning.

### Sensory discrimination training using directional air puffs

The rapid change in stimulus-associated anticipatory licking using air current stimulation in the SAT paradigm suggests that multiwhisker stimulation can be a potent stimulus to drive learning. We next probed the capacity of the animal to discriminate different directions of air currents, using a similar reward schedule as before but where “blank” trials were replaced with an airpuff delivered from a different direction. Training cages were designed so that air currents could be precisely positioned relative to the whiskers, and one direction was selected as the rewarded direction.

Initially we selected a 180 degree difference between the rewarded and unrewarded stimulus, reasoning that this might be the most discriminable stimulus pair. Although the majority of animals showed an increase in L_w_>L_b_ (6/9 animals), on average this difference was not significant at 48 hrs of SAT (Fig 7C-E). Reducing the angular difference between the rewarded and unrewarded stimulus to 15 degrees actually improved mean performance after 48 hrs of SAT (Fig 7F-H). This improvement in performance could be observed regardless of the location of the rewarded direction (either above or below the animal; S1 Fig).

**Figure 7.**
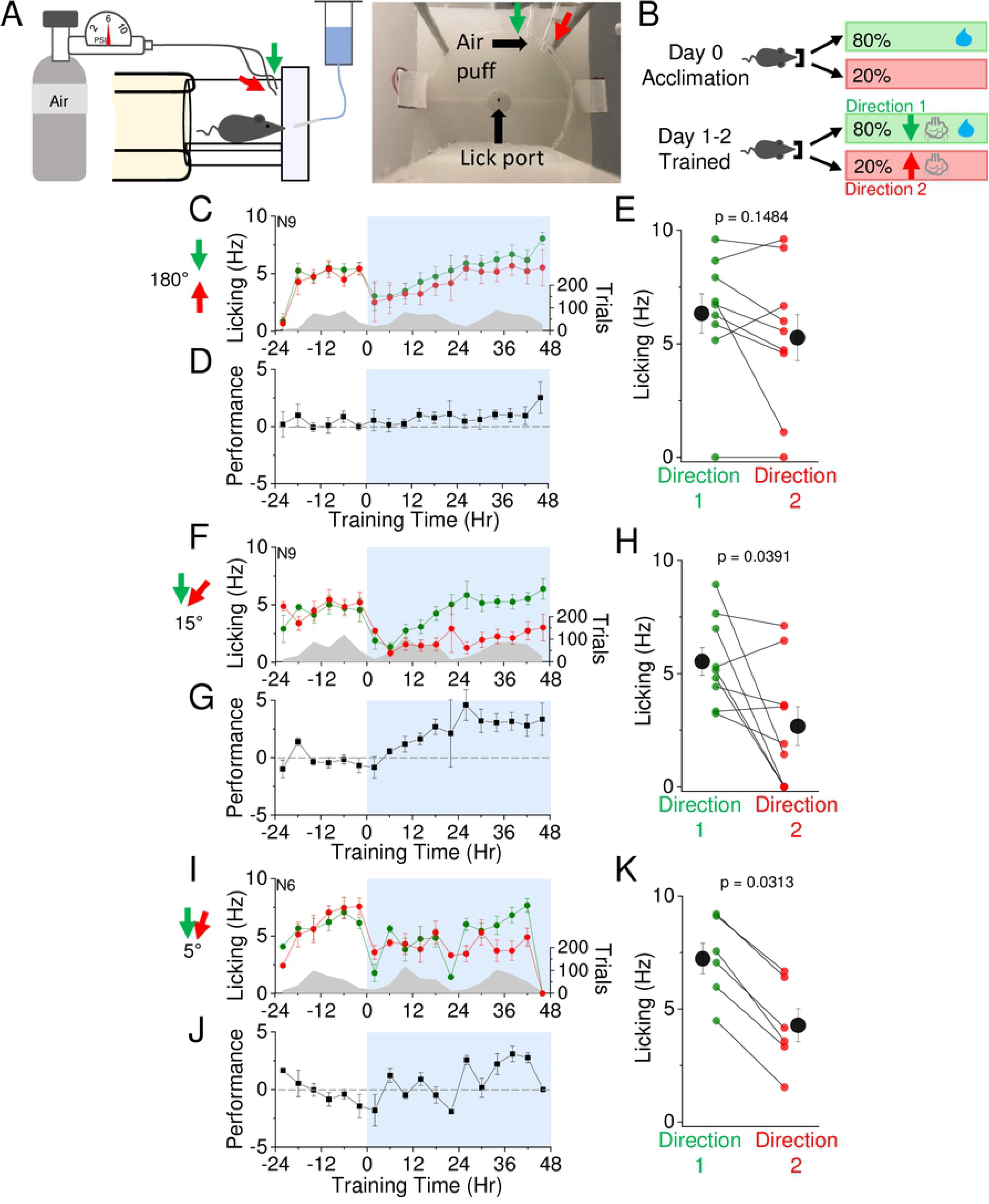
Mice can discriminate air puff direction. A) Left, profile view of bidirectional air puff training apparatus. Right, lick port and air puff position within automated homecage training apparatus. B) Reward contingencies during training. Left, during the initial 24 hour acclimation period, animals receive water on 80% of initiated trials with no air puff. Right, during the training period, animals receive water and an air puff from one direction on 80% of initiated trials, and an air puff from a different direction without water on the remaining 20% of trials. C) Mean lick frequency of animals exposed to air puffs 180 degrees apart for water and blank trials, binned at 4 hr intervals. Air puff association training is indicated at t=0 (12 noon/daylight period). Mean lick frequency for water delivery (green) or blank (red) trials is overlaid upon mean number of initiated trials (grey) across training days. D) Mean performance (lick frequency for water trials – lick frequency for blank trials) for each 4 hour bin during the acclimation period (−24 to 0 hr) and training phases with air puffs 180 degrees apart (0 to 48 hrs). E) Mean lick frequency of each animal exposed to air puffs 180 degrees apart for the last 20% of total trials. N=9 animals. F) Same as in (C) but with animals exposed to air puffs 15 degrees apart, projecting downwards. G) Same as in (D) but with animals exposed to air puffs 15 degrees apart, projecting downwards. H) Same as in (E) but with animals exposed to air puffs 15 degrees apart, projecting downwards. N=9 animals. I) Same as in (C) but with animals exposed to air puffs 5 degrees apart, projecting downwards. J) Same as in (D) but with animals exposed to air puffs 5 degrees apart, projecting downwards. K) Same as in (E) but with animals exposed to air puffs 5 degrees apart, projecting downwards. N=6 animals.

Animals appeared capable of even finer-scale directional discrimination, as further reducing the angular difference between the rewarded and unrewarded direction to 5 degrees continued to show an increase in mean anticipatory licking (Fig 7I-K). These results indicate that mice have an extraordinary ability to differentiate between air current directions and suggest that the facial vibrissae may be specially tuned to this stimulus feature.

### Enhanced reward valence does not improve performance

Changing the water reward to a more desirable reward, such as aspartame, could influence learning trajectory, since mice may initiate more trials in order to obtain more of the desirable reward or because reward signals that regulate learning are stronger. On acclimation day, mice were supplied with water on 80% of initiated nosepokes. At the onset of SAT, water was replaced with an aspartame-containing solution.

Animals showed significant increased anticipatory licking behavior to the stimulus at the end of the 48 hrs training period (Fig 8B-D), similar to the performance of interleaved control animals also trained to the 6 psi stimulus but with a water reward. On average, aspartame-trained animals showed a higher number of trials compared to water-trained animals, a difference that was not significant (mean±SEM: Water 389±85 trials/day, N=6 mice; Aspartame 492±41, N=9 mice; p=0.25). Analysis of behavioral change over time suggested that the increase in licking on stimulus trials versus blank trials for aspartame-trained animals might be delayed compared to mice that only received water. This delayed separation suggests that at least in our experimental set-up, aspartame replacement does not facilitate learning trajectory. These findings are consistent with prior results indicating that reward palatability does not strongly influence learning in rodents (16).

**Figure 8.**
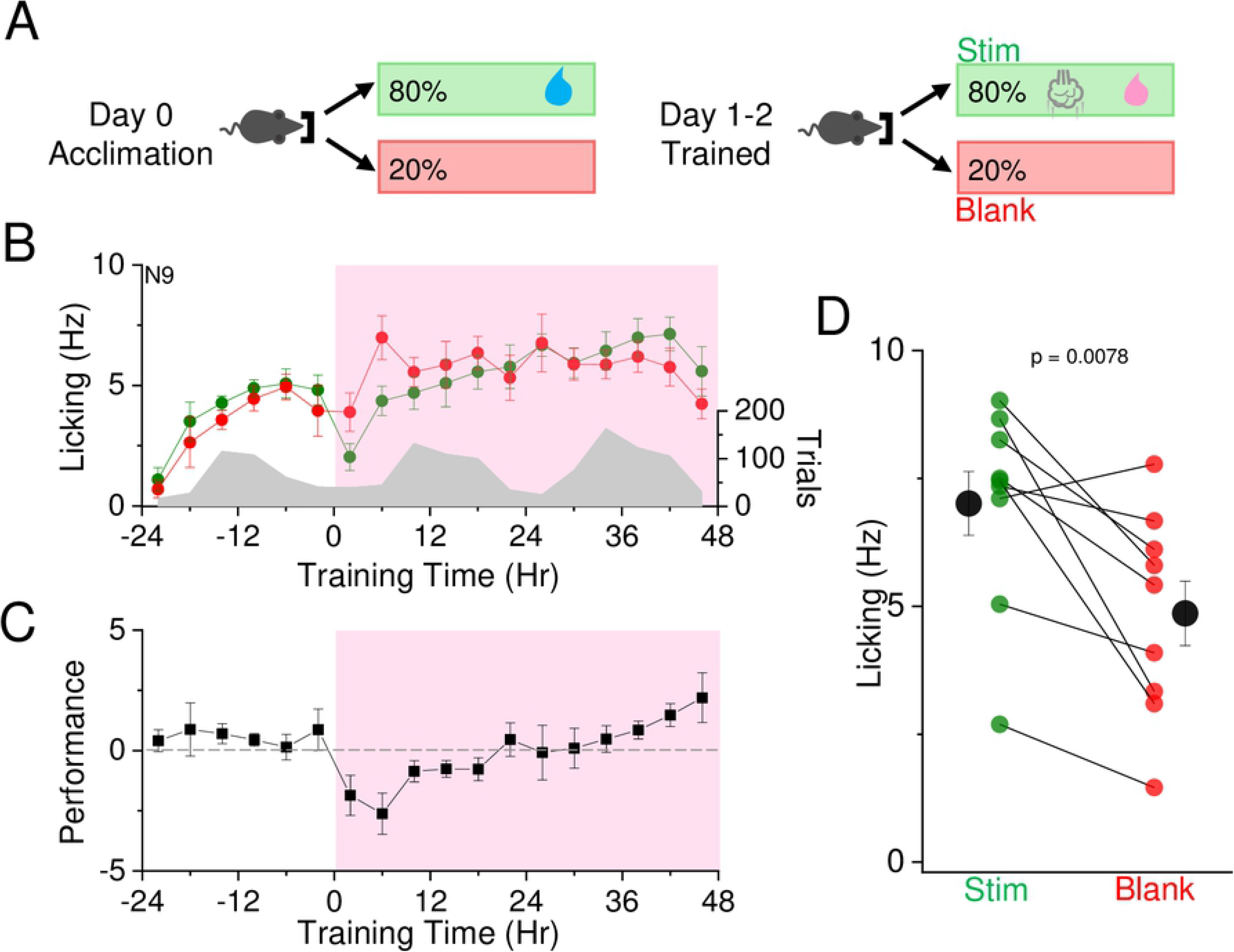
Aspartame does not enhance learning with 48 hrs SAT. A) Reward contingencies during training. Left, during the initial 24 hour acclimation period, animals receive water on 80% of initiated trials with no air puff. Right, during the training period, animals receive air puff and aspartame on 80% of initiated trials. B) Mean lick frequency for water and blank trials, binned at 4 hr intervals. Air puff set at 6 psi and association training is indicated at t=0 (12 noon/daylight period). Mean lick frequency for water/aspartame delivery (green) or blank (red) trials is overlaid upon mean number of initiated trials (grey) across training days. C) Mean performance (lick frequency for water/aspartame trials – lick frequency for blank trials) for each 4 hour bin during the acclimation period (−24 to 0 hr) and training phases (0 to 48 hrs). D) Mean lick frequency for the last 20% of total trials for each animal exposed to aspartame and 6 psi air puff. N=9 animals.

## Discussion

### Summary

We developed a homecage, automated behavioral training system for freely-moving animals that coupled a multiwhisker stimulus with water reward. We extended our previous study (11) by modulating training parameters to investigate the whisker-dependence, stimulus-intensity, reward frequency and valence, and directional discrimination capabilities of animals trained in this environment. We find that multiwhisker stimulation is a potent sensory modality to drive sensory learning, and our results establish that multiwhisker stimulation is a robust and easily adapted system for sensory association training in mice.

### Benefits of automated training

Automated behavioral paradigms, such as the IntelliCage (17–20), have been used in other studies to train animals for discrimination of odors (21–23), oriented lines (24,25), and auditory tones (26–28). Several studies also used automated set-ups for motor control tasks (29–32). Automated training improves standardization across experimenters and different laboratories (32,33) and allows for minimal experimenter contact with mice which reduces the stress associated with handling (34–36). In addition, the use of a freely-moving behavioral paradigm carried out in the homecage environment further reduces stress (33–35). In our assay, animals were not water restricted, an arrangement that further reduces animal stress and the burden of experimenter monitoring and documentation (37).

Automated training significantly increases experimental throughput, a particular advantage for experiments that require large numbers of animals (11,27). Importantly, behavioral data were automatically collected using our custom-designed Arduino system, allowing remote data access and rapid analysis for long training periods. Due to this ease of use, our automated sensory training system could be used for phenotypic characterization of mutant strains (see for example (38–42)).

We designed an automated behavioral training chamber that would reliably deliver a multiwhisker stimulus. An advantage of multiwhisker stimulation is that it can be delivered without the application of artificial agents for magnetic whisker deflection (43,44), does not require precise animal positioning for delivery and is thus suitable for freely-moving animals, and provides a large anatomical area – >400 um^2^ of the posterior-medial barrel subfield representing the large facial vibrissae –for detailed anatomical and neurophysiological analysis.

### Task design and classical conditioning

Although SAT in this study has many components of a classical Pavlovian conditioning task, it differs in several important respects. First, because animals self-initiate trials, there is an operant aspect to the trial design. Second, the response to the stimulus, licking, is under voluntary control (45). Furthermore, the learned behavior contains the incentive of receiving water which is aligned with operant conditioning since Pavlovian conditioning has no incentives associated.

The 500ms delay between air puff termination and water delivery in this task classifies it as a trace conditioning. This type of conditioning is different than delay conditioning in which one stimulus is presented, followed by a second stimulus, and both stimuli are then terminated at the same time, a training paradigm that may engage different neural circuits in the brain. Delay conditioning typically requires the cerebellum and is not associated with a conscious awareness of the relationship between the two stimuli. Trace conditioning also requires the cerebellum, however, the hippocampus and neocortex are additionally needed for accurate completion of the task (46). Therefore, SAT as implemented by this homecage training environment is likely to engage multiple brain circuits and may be well-suited for the analysis of cortical circuit changes during learning.

### Training modifications using multiwhisker stimulation

An advantage of this training set up is that multiwhisker stimulation parameters can be adjusted for a large variety of learning objective. For example, animals can be trained to detect whisker stimulation one side of the face, and then tested on association learning with stimulation to the opposite side. Different patterns (duration, frequencies, or directions) of multiwhisker stimuli can be used to probe more complex forms of associative learning. In addition, water delivery can be decoupled from the stimulus in this training environment to look at stimulus-dependent changes in cortical response properties in the absence of learning (11).

### Animal-to-animal variability

Using a multiwhisker stimulus to drive associative learning behavior revealed a substantial amount of variability across animals in learning trajectories in our study. This variability was captured by reported raw values for lick rates, instead of a d’ measurement that normalizes behavioral measurements. What might account for this? Using a simple criterion of greater lick frequency in stimulus versus blank trials (L_w_>L_b_) as evidence of learning, we observed that weak airpuff intensity was correlated with a marked reduction in the fraction of animals that learned. At 48 hrs of SAT with 1-2 psi, 55% of mice showed L_w_>L_b_, but with 9 psi, 86% of animals showed this. It was clear that the 9 psi stimulus was more salient, as more than half the animals stopped approaching the lickport for water in this condition, likely due to the aversive quality of the high-intensity airpuff. At 1-2 psi, all animals participated throughout the training period, consistent with the interpretation that lower stimulus intensities may be less aversive but may be more difficult to detect, particularly for some animals. Differences between animal strategies for receiving the stimulus may also explain across animal heterogeneity in performance (see for example (47)). Although our automated approach sought to reduce animal stress from handling that can influence learning behaviors in mice (48), individual mice can show variable levels of anxiety that can also influence learning (49). The reported variability in behavioral performance during this automated SAT paradigm may be useful in examining causal relationships between learning and cellular and synaptic changes in the mouse brain.

## Acknowledgements

Special thanks to Dylan McCreary, Rogan Grant, and Stefan Bernhard for early contributions to cage design, and members of the Barth lab for critical comments on the manuscript.

## Author Contributions

Conceptualization: Sarah Bernhard, Jiseok Lee, Andrew Hires, Alison Barth

Formal analysis: Sarah Bernhard, Alex Lee, Andrew Erskine

Funding acquisition: Sarah Bernhard, Sam Hires, Alison Barth

Investigation: Sarah Bernhard, Jiseok Lee, Mo Zhu

Methodology: Sarah Bernhard, Alex Lee, Andrew Erskine, Sam Hires,

Validation: Sarah Bernhard, Jiseok Lee, Mo Zhu

Visualization: Sarah Bernhard, Alison Barth

Writing ± original draft: Alison Barth

Writing ± review & editing: Sarah Bernhard and Alison Barth

## Supporting information

**S1 Video. Example video clip taken for whisker video analysis at 1 psi**

**S2 Video. Example video clip taken for whisker video analysis at 5 psi**

**S3 Video. Example video clip taken for whisker video analysis at 9 psi**

**S1 Figure. Discrimination using directional air puff alters learning rate at 15 degrees.** A) Profile view of bidirectional air puff training apparatus. B) Reward contingencies during training. Left, during the initial 24 hour acclimation period, animals receive water on 80% of initiated trials with no air puff. Right, during the training period, animals receive water and an air puff from one direction on 80% of initiated trials, and an air puff from a different direction without water on the remaining 20% of trials. C) Mean lick frequency of animals exposed to air puffs 15 degrees apart for water and blank trials, binned at 4 hr intervals. Air puff association training is indicated at t=0 (12 noon/daylight period). Mean lick frequency for water delivery (green) or blank (red) trials is overlaid upon mean number of initiated trials (grey) across training days. D) Mean performance (lick frequency for water trials – lick frequency for blank trials) for each 4 hour bin during the acclimation period (−24 to 0 hr) and training phases with air puffs 15 degrees apart (0 to 48 hrs). E) Mean lick frequency of each animal exposed to air puffs 15 degrees apart for the last 20% of total trials. N=7 animals.

### Suggested Reviewers

Jerry Chen

David Margolis

Aric Agmon

Tansu Celikel

Dan O’Connor

